# Multidecadal decline and recovery of aspen experiencing contrasting fire regimes: long-unburned, infrequent and frequent mixed-severity wildfire

**DOI:** 10.1101/2020.04.28.065896

**Authors:** Cerena J. Brewen, John-Pascal Berrill, Martin W. Ritchie, Kevin Boston, Christa M. Dagley, Bobette Jones, Michelle Coppoletta, Coye L. Burnett

## Abstract

Quaking aspen (*Populus tremuloides*) is a valued, minor component on western landscapes. It provides a wide range of ecosystem services and has been in decline throughout the arid west for the last century. This decline may be explained partially by the lack of fire on the landscape as aspen benefit from fire that eliminates conifer competition and stimulates reproduction through root suckering. Managers are interested in aspen restoration but there is a lack of knowledge about their spatial dynamics in response to fire. Our study area in northeastern California on the Lassen, Modoc and Plumas National Forests has experienced recent large mixed-severity wildfires where aspen was present, providing an opportunity to study the re-introduction of fire. We observed two time periods; a 54-year absence of fire from 1941 to 1993 preceding a 24-year period of wildfire activity from 1993 to 2017. We utilized aerial photos to delineate aspen stand size, location and succession to conifers. We chose aspen stands in areas where wildfires overlapped (twice-burned), where only a single wildfire burned, and areas that did not burn within the recent 24-year period. We looked at these same stands within the first period of fire exclusion for comparison (i.e., 1941-1993). In the absence of fire, all aspen stand areas declined and all stands experienced increases in conifer composition. After wildfire, stands that burned experienced a release from conifer competition and increased in stand area. Stands that burned twice or at high severity experienced a larger removal of conifer competition than stands that burned once at low severity, promoting aspen recovery and expansion. Stands with less edge:area ratio also expanded more with fire present. Across both time periods, stand movement, where aspen stand footprints were mostly in new areas compared to footprints of previous years, was highest in smaller stands. In the fire exclusion period, smaller stands exhibited greater changes in area and location (movement), highlighting their vulnerability to loss in the absence of disturbances that provide adequate growing space for aspen over time.

## Introduction

From the mid-1800s, changing land-use patterns and active wildfire suppression throughout the western United States has altered fire regimes [1, 2, 3, 4]. In frequent-fire forest types, wildfires historically maintained more open forest conditions [5, 1, 6]. In the absence of wildfire disturbances over much of the 20^th^ Century, stand density has increased steadily [7]. These conditions have promoted a shift in species composition towards more shade-tolerant species [8], and are not favorable for light-demanding pioneer species such as aspen [9, 10, 11]. Aspen have declined 50-90% in the western United States due to drought, insects and disease, ungulate browsing, and lack of disturbance [11,12]. Absence of fire may be contributing to aspen decline, as fire is an important disturbance agent promoting persistence of aspen in many stands [10, 13, 14].

Quaking aspen (*Populus tremuloides*) is a valuable, although infrequently encountered, component on northeastern California landscapes [12]. Aspen ecosystem services range from biodiversity hotspots of plant and animal species, economic value, cultural significance and aesthetic appeal [15, 16]. The vegetation composition associated with aspen stands is richer than other ecotypes on a shared landscape [15]. As this rich vegetation promotes higher volumes of insects, more bird species have been found in aspen stands than in neighboring conifer stands due to this greater food source [17, 18]. Bat species have also been found to select aspen cavities over conifer cavities due to cooler temperatures in aspen trees [19]. Additionally, aspen stands have been observed as small mammal hotspots within conifer dominated landscapes [20].

Aspen regenerates primarily through root suckers and less commonly through seed [9]. Because of this, once aspen is extirpated from a site, it is difficult for the species to become re-established. Other tree species that can grow in association with aspen, such as pines and firs that are long-lived and reproduce by seed more reliably than aspen, will most likely replace aspen [21]. In the Sierra Nevada mountains of California and Nevada, aspen functional types include either pure aspen stands that appear stable or seral aspen coexisting in mixture with conifers; these are vulnerable to replacement by conifers in the absence of disturbance [22]. Disturbances such as timber harvesting and fire promote aspen regeneration [23]. After introduced fire, aspen were observed to almost double their sucker counts from preburn conditions [24]. After mixed-severity wildfires, aspen patches that burned at high severity exhibited the greatest density of aspen root suckering [25, 26, 27]. High densities of young aspen are necessary when high herbivory pressure is present, allowing new aspen cohorts to establish and preventing potential loss of aspen from overgrazing. High-severity wildfire is also important for relieving pressures of competition. After trees are killed by high severity fire, aspen root suckers quickly re-occupy growing space liberated by the disturbance. In the absence of disturbance, increasing stand density and competition reduces growth and vigor of aspen and increases competition induced mortality [28]. Under these conditions, aspen continue to produce root suckers, but they rarely reach sapling sizes and are therefore unlikely to recruit to the overstory [29].

Aspen responds well to compound disturbances, including when another disturbance precedes fire (e.g., insect outbreaks, disease and wind storms) [30]. While non-sprouting species may be killed and lose their seed source over the course of multiple disturbances, we expect aspen to regenerate by root suckering in response to each disturbance and progressively increase occupancy relative to competing conifers. After a single fire, aspen regenerate rapidly, but other fire-adapted tree species, such as lodgepole pine (*Pinus contorta*), may also regenerate well. [30]. Positive regeneration responses to wildfire and to compound disturbances suggest aspen may benefit from frequent wildfire or managed fire events. Wildland fire use is a management tool where naturally ignited fires are allowed to burn in areas based on cultural or natural resource objectives. When wildland fire use was initiated in the Illilouette Creek Basin in Yosemite National Park in 1990, there were many areas that reburned, showing that overlap of managed fire is possible [31]. It was also shown that wildland fire use can result in burning that is very similar to historical fires (in terms of severity), even after a history of fire exclusion [32]. Managing wildfires may be the most effective method of promoting regeneration and vigor of declining, fire-adapted species such as aspen.

Aspen are threatened by drought [35], climate change [36], and succession to conifer [37]. As conifers proceed to replace aspen throughout the west [12], aspen may be restricted to less favorable sites. Conversely, disturbances may let aspen recolonize areas of higher site quality. There is limited understanding of the potential for fire to kill conifers and concomitantly allow aspen stands to expand in area or move into new areas. The Sierra Nevada mountain range and adjacent areas of northern California continue to experience different extents, severities and overlapping of naturally occurring fires [33]. However, within the same region, aspen has also experienced up to 24% loss in the South Warner Mountains within 50 years [34]. This suggests a need for studies of how individual aspen stands have changed throughout time, especially their response to disturbance or lack thereof [38].

Observations of historical conditions are required to study long-term changes in aspen stands. A combination of current satellite imagery and historic aerial photos provide a time series of imagery that allow us to observe change in tree species composition and extent of aspen stands. Landsat spatial data is a preferred satellite imagery source as it was designed to detect long-term patterns of change on Earth’s landscapes [39]. Landsat digital imagery can be used for tree species detection to an accuracy of 88.8% consistency of ground truth data [40]. Landsat has been used for measuring temporal change in species composition and stand extent by using imagery from multiple dates with an accuracy of up to 87% [41]. Google Earth is a source of Landsat imagery that provides recent and past imagery over 25 years in some areas [42]. When images that predate satellite imagery technology are needed, aerial photography may be available. Determination of species composition and single tree analysis has been accomplished through the use of aerial photos with an accuracy of 70-80% [43].

Studies featuring long-term observations of aspen stand spatial dynamics are few and none have looked at the effect of overlapping wildfire. Identifying factors related to aspen decline and expansion would help to inform land managers seeking to increase aspen area on the landscape. Our study objectives were to compare aspen stand area change, movement and succession to conifers among stands that have experienced different fire frequencies and severities. Our 76-year study period started with a 52-year absence of fire to observe stand conditions without disturbance, followed by a 24-year period where wildfire activity was present with stands experiencing three different fire regimes:

A. frequent fire (2 overlapping fire footprints within the last 24 years),
B. infrequent fire (1 fire within the last 24 years) and
C. fire exclusion (absence of fire within the last 76 years).

We hypothesized that spatial dynamics (expansion/contraction of stand area and movement of stand boundaries) were driven by wildfire activity. Specifically, we expected: 1) stand area to recede and move in the absence of fire, 2) associated increases in conifer dominance in the absence of fire, and 3) greater stand expansion in the presence of frequent wildfires than a single wildfire event.

## Methods

### Project Area

The study sites are located on US Forest Service lands in northeastern California on the Lassen, Modoc and Plumas National Forests (Fig 1). Aspen stands were selected for analysis within large fire footprints on each forest: the 2000 Storrie fire (Lassen NF), the 2001 Blue complex (Modoc NF) and the 2007 Moonlight fire and Antelope complex (Plumas NF). Within each main fire footprint were areas where other fires had burned within the 24-year study period and aspen was present. The 2012 Chips fire reburned areas within the Storrie fire footprint, the 1994 Corporation fire footprint was partly reburned by the Blue complex footprint, and the 2001 Stream fire footprint was partly reburned by the Moonlight fire and the Antelope complex.

**Fig 1.**
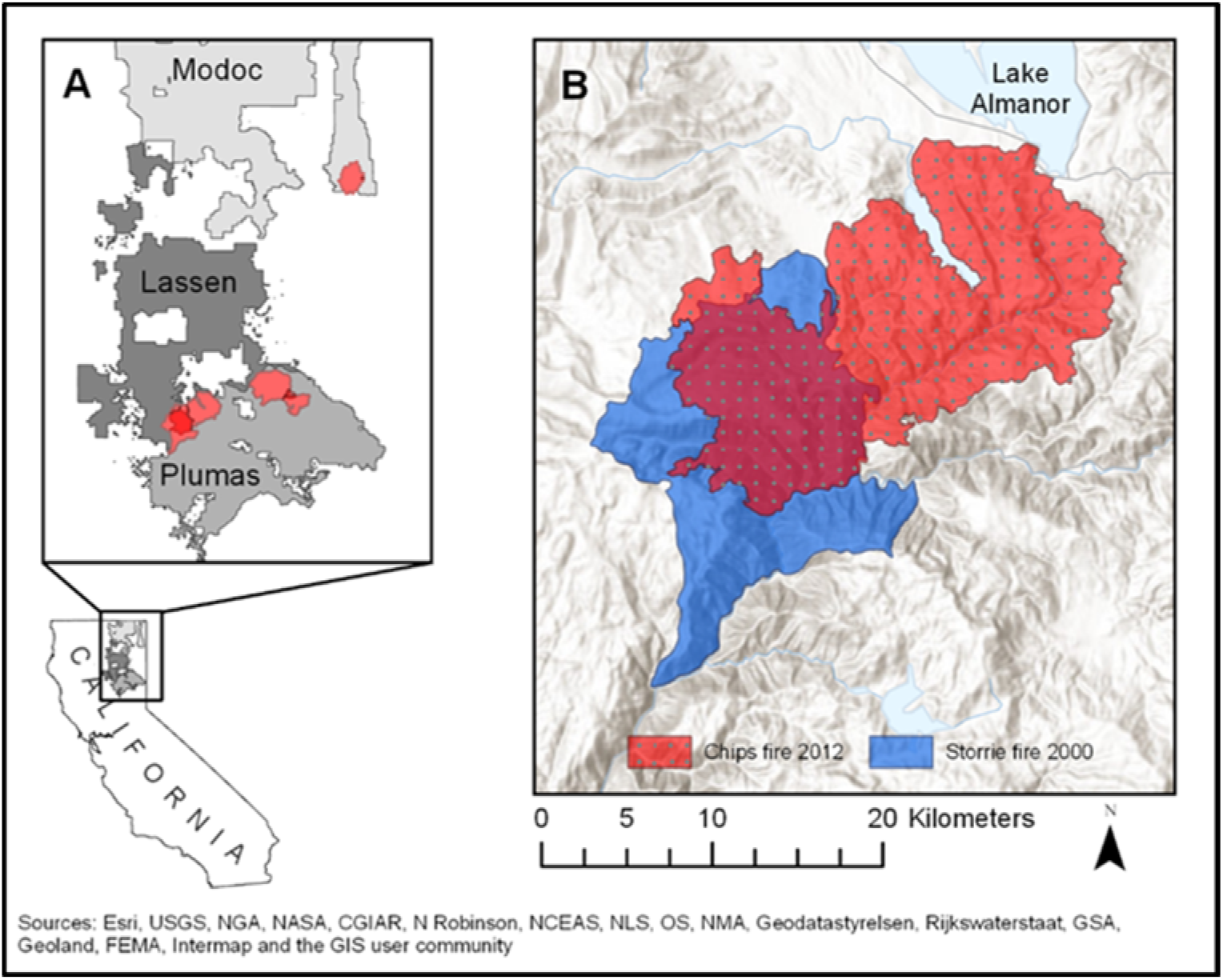
A) Location of study wildfires on three National Forests in northeastern California, and B) example area of wildfire overlap on the Lassen National Forest.

Aspen stands within the overlapping fire footprints represent the ‘frequent fire’ condition where two fire disturbances were experienced within 24 years. Stands representing only one instance of ‘infrequent fire’ were selected within the main fire footprint only. A similar number of unburned stands were chosen from outside fire footprints but as close as possible to burned stands in an attempt to minimize site differences among burned and unburned aspen stands.

### Aspen Stand Measurements

We completed field assessments of our 33 study stands between 2014 and 2018. Stand boundaries were mapped with a resource-grade GPS unit and stand details were documented, including the number of cohorts present, estimates of conifer encroachment, and presence of insects and/or diseases (Table 1). Mapped polygons of aspen from the stand assessments were overlaid on aerial images from past years as a guide to help identify and delineate aspen in each historical image. New polygons were created around aspen visible in each image. Polygons in recent years (1993-2017) were digitized on aerial images in Google Earth. Historic (1941) extent of aspen visible from aerial images was digitized on georeferenced aerial photos from that year. Aerial photos from 1941 on the Lassen and Plumas National Forests were previously georeferenced by US Forest Service employees. Photos of the Modoc National Forest were scanned at 600dpi and georeferenced in ArcMap [44].

**Table 1.** Number of aspen stands by disturbance type, and range of stand areas.

| Forest | n | Unburned | 1 Burn | 2 Burns | Min (ha) | Max (ha) | Mean (ha) |
| --- | --- | --- | --- | --- | --- | --- | --- |
| Lassen | 10 | 4 | 3 | 3 | 0.02 | 1.08 | 0.29 |
| Modoc | 9 | 2 | 5 | 2 | 0.02 | 5.21 | 1.63 |
| Plumas | 14 | 3 | 6 | 5 | 0.09 | 8.84 | 1.45 |

All polygons from each year were brought together on one map file in ArcMap for each of the three forests. Within the attribute table of each year, all polygons were assigned a stand number based on site type and proximity to other polygons (Fig 2).

**Fig 2.**
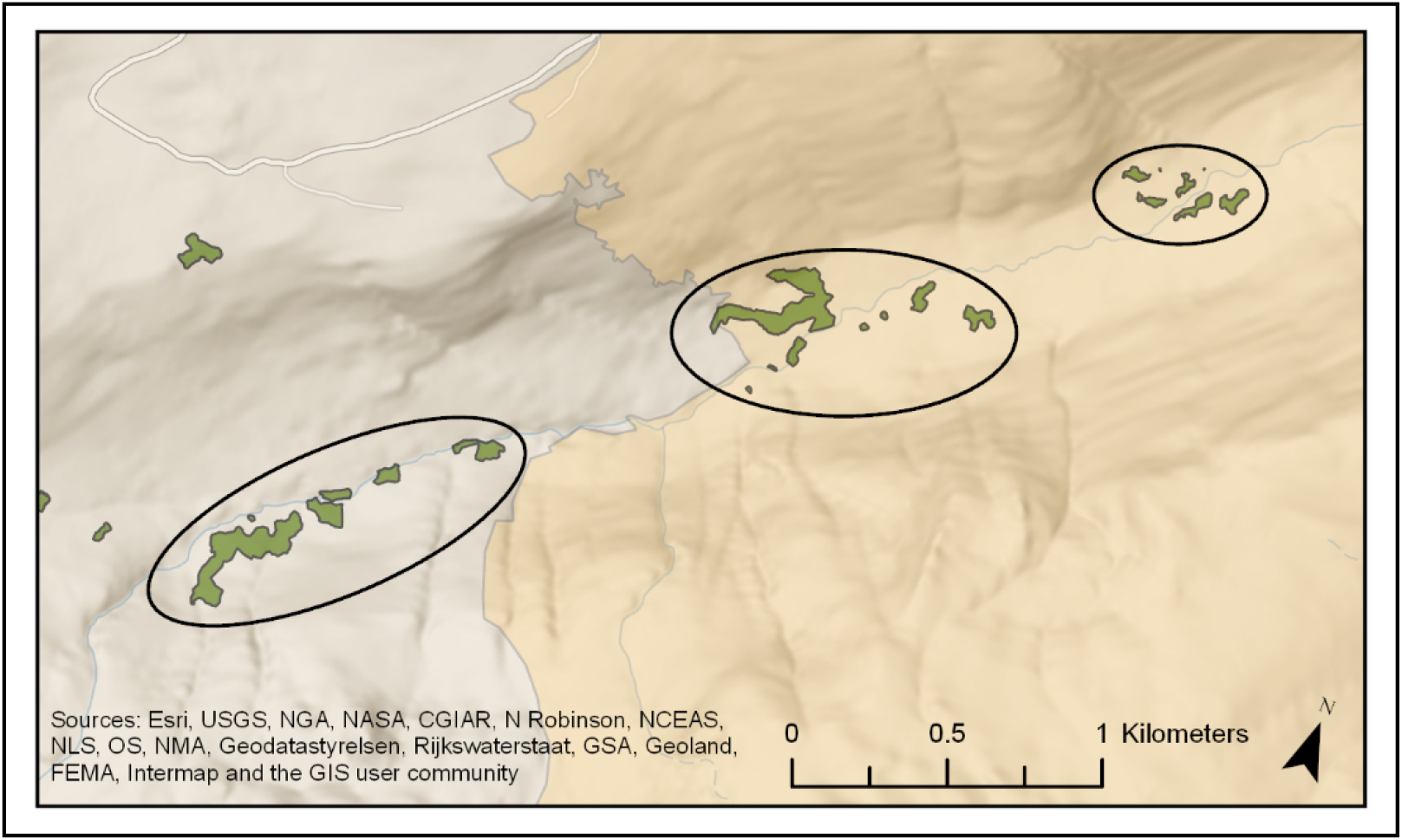
Example of aspen polygon grouping procedure where all polygons in close proximity (shown here within an ellipse) were grouped together as a single stand.

### Variable Calculations

To evaluate change in aspen over time, we derived the following three response variables: proportional area change (A_1_/A_0_) as a ratio; movement of stand boundaries; and change in conifer cover percent. We also derived the following predictor variables: edge:area ratio, fire severity, and a composite fire severity-frequency score.

To estimate proportional area change, we calculated the total area in hectares for all stands at each time period in ArcMap. Proportional change was calculated for two different time periods: the fire exclusion period (1941-1993) and the return of fire period (1993-2017). Periodic proportional change was calculated by dividing the area at the end of the period with the area at the beginning: Δ = A_1_/A_0_ for both time periods. There were a few stands that were undetectable in the 1993 aerial imagery but visible in 2017 after fire, indicating that these stands were not completely lost in 1993. In these stands, we assumed that at least one individual was present prior to 2017, and assigned an area of 0.003 hectares in 1993, which is the size of a patch that only includes a single tree, representing the smallest measurable area.

Stand movement was calculated using the union tool to combine polygons from all three years into one shapefile. This gave us new polygons for areas lost between assessment years, areas gained between years and areas maintained between years for each aspen stand. We then used the calculate geometry function again to get area values for each of these polygons. Stand movement values were calculated as the proportion of a recent year’s stand area that was “new” because it was found outside of the previous assessment year’s stand area footprint:

M_1941-1993_ = 1993new_area/1993total_area
M_1993-2017_ = 2017new_area/2017total_area

For example, a stand that remained entirely within the prior year’s footprint would have movement value = 0; a stand that occupied an entirely new location completely outside the prior year’s footprint would have movement value = 1 (i.e., maximum movement).

Percent conifer composition was measured using ocular estimates in the imagery from each year. Conifer composition in 1993 and 2017 was observed on imagery in Google Earth and 1941 conifer composition was observed on aerial photos displayed in ArcMap. For each year, the field-derived GPS polygons from the recent field assessments of aspen stand area were overlaid on that year’s imagery, along with the digitized stand delineation polygon from that year to account for areas where aspen was observed during ground assessments but not from aerial imagery. A number between zero and 100 was assigned to each stand in each year based on the conifer crown extent as a percentage of stand area observed within the field-derived and the digitized polygons. Conifer composition change was then calculated by subtracting the percentage for the ending year’s conifer composition from the percentage for the starting year’s conifer composition for each time period, resulting in a positive number for conifer composition increase and a negative number for conifer composition decrease (e.g., increasing conifer: 50% - 10% = 40% increase; declining conifer 20% - 35% = −15%).

Stand edge:area ratio was derived for each time period in ArcMap. Edge:area ratio was determined by calculating the perimeter (meters) of each stand and dividing it by total stand area (square meters).

Fire severity was assigned to each stand using Google Earth. In and around each stand, conifer mortality was assessed by comparing conifer cover in years before and after all fire events. Four fire severity categories were assigned (unburned, low, moderate, high), where low severity had little to no conifer mortality, moderate severity had visible patches of conifer mortality and high severity had 100% conifer mortality [45]. Severity in stands that burned twice was measured as a cumulative effect, as separate severity for each fire was unobservable. For statistical analysis, low and moderate stands were grouped together as there was only one stand that burned at low severity.

Fire frequency (FF) and fire severity (FS) were combined into a single categorical variable due to singularity: FFFS. Five categories were constructed: unburned, once-burned experiencing low/moderate severity, once-burned experiencing high severity, twice-burned experiencing low/moderate severity and twice-burned experiencing high severity.

### Spatial Dynamics Modeling

A t-test in R software version 3.5.2 [46] was used to test for a fire frequency effect on proportional area change when fire was present between years 1993 and 2017. The t-test compared proportional area change in once versus twice-burned stands.

Linear models were constructed to test for variables influencing three response variables (i) proportional stand area change, (ii) proportional stand movement, and (iii) conifer composition change in each time period. Proportional area change data for the return of fire period were logarithmically transformed to reduce skewness. The lm function was used to model proportional area change. Stand movement was modeled using the betareg function, accounting for data confined between zero and one. Stand movement values of 1 were changed to 0.999999 to use betareg analysis. Two Modoc aspen stands were not included in the stand movement models between 1941 and 1993 due to georeferencing difficulties, leaving 31 stands available for stand movement analysis. The glm function was used to model conifer composition change as a function of candidate explanatory variables (Table 2). Fire severity was not included as a candidate variable for the conifer composition change models as conifer composition before and after fires was used to measure fire severity, making these two variables inherently related. Therefore, fire frequency alone was tested in conifer composition change models.

**Table 2.**
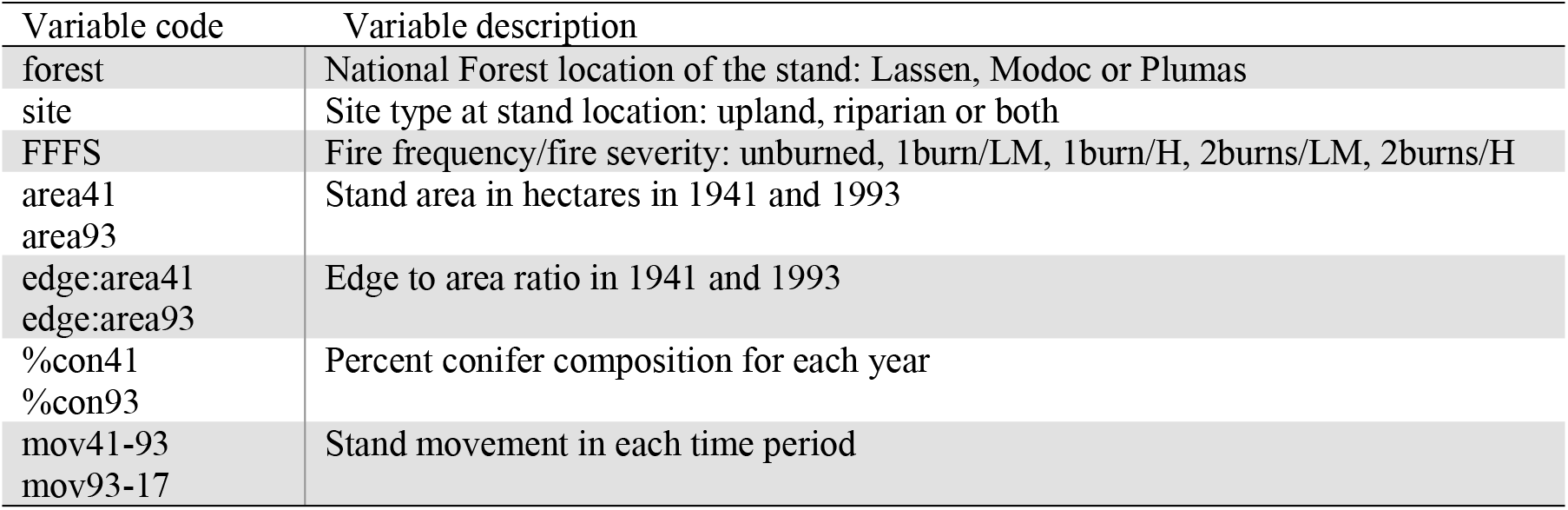
List of candidate explanatory variables for analysis of changes in aspen-conifer stands. Fire severity within the FFFS category are coded as LM for low/moderate severity and H for high severity.

| Variable code | Variable description |
| --- | --- |
| forest | National Forest location of the stand: Lassen, Modoc or Plumas |
| site | Site type at stand location: upland, riparian or both |
| FFFS | Fire frequency/fire severity: unburned, 1burn/LM, 1burn/H, 2burns/LM, 2burns/H |
| area41<br>area93 | Stand area in hectares in 1941 and 1993 |
| edge:area41<br>edge:area93 | Edge to area ratio in 1941 and 1993 |
| %con41<br>%con93 | Percent conifer composition for each year |
| mov41-93<br>mov93-17 | Stand movement in each time period |

Backwards eliminations were used for selection of variables used in best models as determined by Akaike Information Criterion corrected for small samples (AICc) [47]. Pairwise comparisons were performed on categorical variables to determine significant differences among individual categories using the eemeans package in R [48]. The coefficients generated from the pairwise comparisons were used to represent least-squares mean and standard error values for each category, and plotted to depict differences among categories of each categorical variable within the best models. Expected values across the range of observed values for each continuous variable were also plotted using the ggplot2 package in R [49].

## Results

On average, our study aspen stands were much larger and had fewer conifers in 1941 than in 1993 (Table 3). Over the 76-year study period (1941-2017), we observed an overall decline in aspen stand area where fire was absent. More recently, however, stands experiencing fire collectively exhibited area expansion. Changes in total aspen stand area (ha) within each fire frequency category were different from each other (p = 0.0451) when fire was present between years 1993 and 2017 (Fig 3). Individual stands exhibited variability in how their area changed with and without wildfire disturbances (Fig 4). Stands movement varied widely, ranging from zero movement to full movement into completely different locations. Conifer composition was also highly variable in space and time, but generally increased from 1941-1993 and decreased from 1993-2017 (Table 3).

**Table 3.** Summary data for continuous response and predictor variables.

|  |  | Min. | Max. | Mean |
| --- | --- | --- | --- | --- |
| *Response Variable* | Proportional area change: 1941 - 1993 | 0.00 | 0.69 | 0.23 |
|  | Proportional area change: 1993 - 2017 | 0.00 | 14.70 | 2.64 |
|  | Stand movement: 1941 -1993 | 0.00 | 1.00 | 0.65 |
|  | Stand movement: 1993 - 2017 | 0.17 | 1.00 | 0.74 |
|  | Conifer comp. change: 1941 - 1993 | -0.27 | 0.92 | 0.29 |
|  | Conifer comp. change: 1993 - 2017 | -0.90 | 0.18 | -0.29 |
| *Predictor Variable* | area41 | 0.08 | 11.11 | 2.49 |
|  | area93 | 0.00 | 6.67 | 0.73 |
|  | edge:area41 | 0.02 | 0.19 | 0.08 |
|  | edge:area93 | 0.00 | 0.64 | 0.18 |
|  | %con41 | 0.02 | 0.65 | 0.25 |
|  | %con93 | 0.05 | 1.00 | 0.53 |

**Fig 3.**
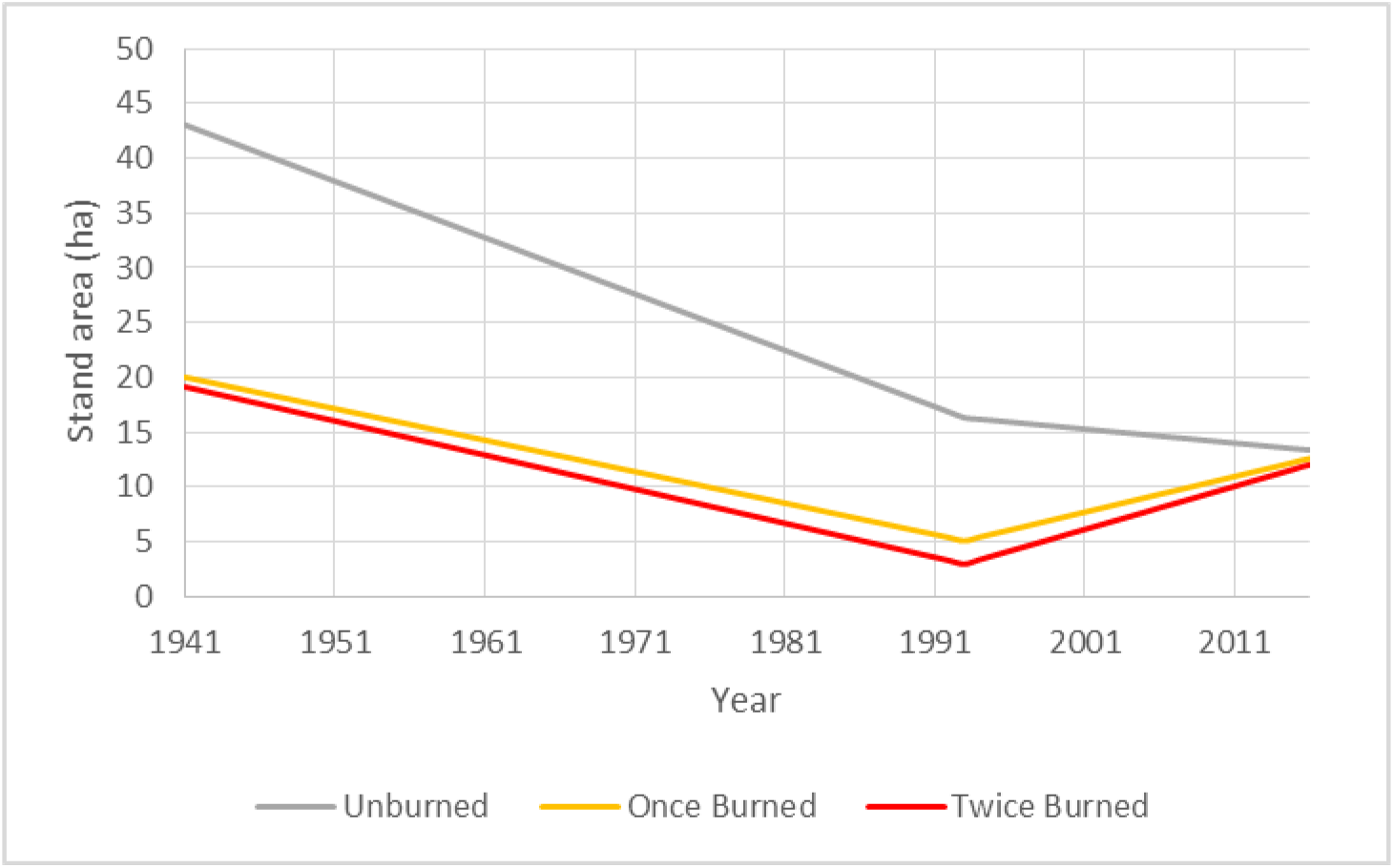
Total aspen stand area in each fire frequency category in 1941, 1993, and 2017, connected by lines depicting average annual change.

**Fig 4.**
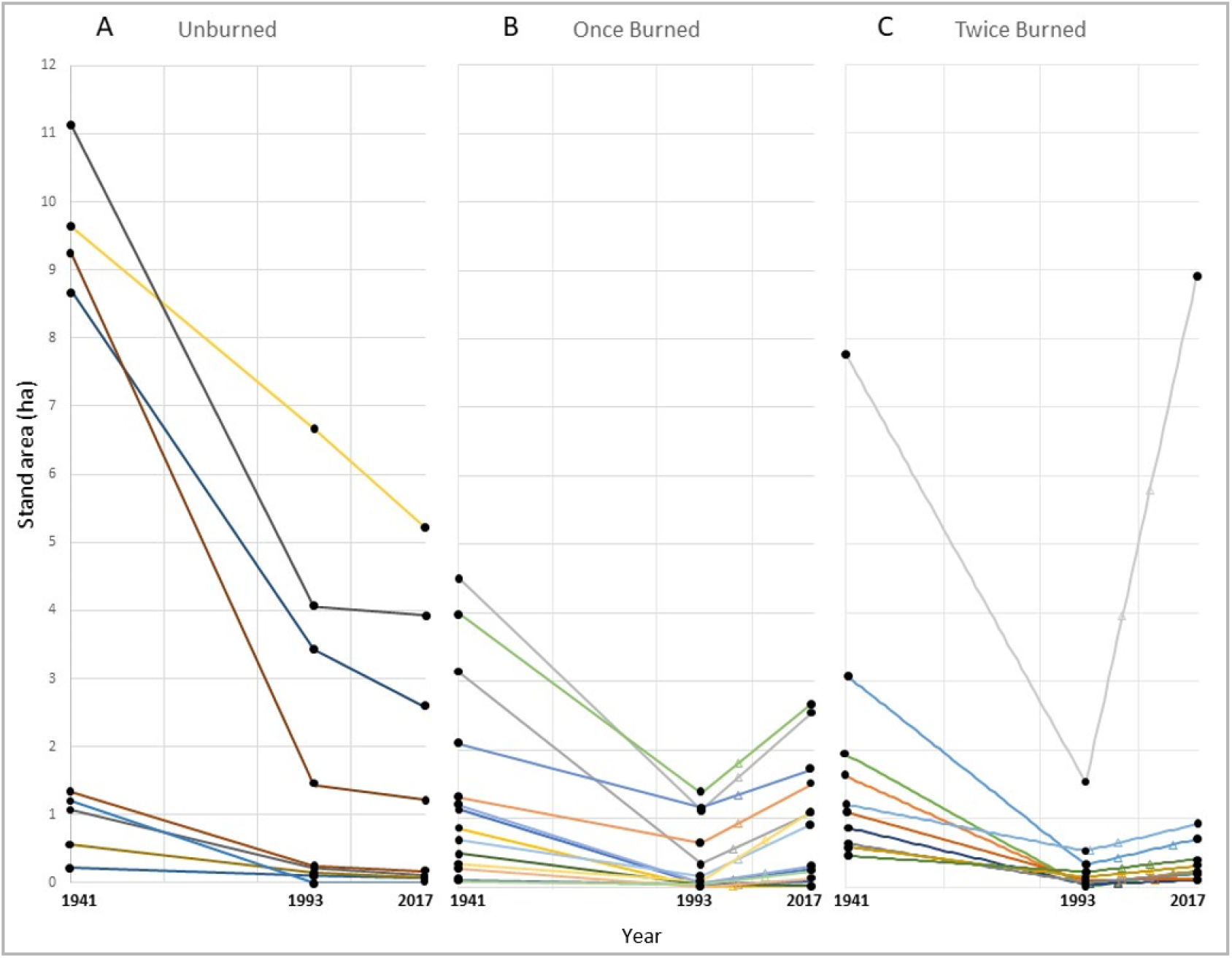
Stand area change over time for individual aspen stands in each fire frequency category: unburned (A), once-burned (B) and twice-burned (C). Triangles denote years where a stand burned between 1993 and 2017.

### Period of Fire Exclusion (1941 – 1993)

The period of fire exclusion was characterized by decline in area of all aspen stands at all of our study sites. Rate of area decline was greater among stands that had lower edge:area ratios in 1941 (p = 0.0003) and smaller stand areas in 1941 (p = 0.0007). Aspen stands on the Lassen declined most rapidly, followed by the Plumas and then the Modoc (Fig 5, Table 4).

**Table 4.** Model summaries for selected models in the period of fire exclusion (1941-1993) for 33 stands in the proportional area change and conifer composition change models and 31 stands in the stand movement model. Variance inflation factor (VIF) quantifies multicollinearity, where a value of 10 or above indicates influential multicollinearity [50].

| Response | Parameter | Coeff | S.E. | Pr(>\|t\|) | VIF |
| --- | --- | --- | --- | --- | --- |
| Proportional area change<br>(Adjusted $R^2 = 0.41$) | Intercept | -0.26196 | 0.110 | 0.0197 | |
|  | Modoc | 0.29020 | 0.080 | 0.0007 | 1.5 |
|  | Plumas | 0.04730 | 0.060 | 0.4641 | 1.5 |
|  | edge:area41 | 3.46814 | 0.847 | 0.0003 | 2.3 |
|  | area41 | 0.04423 | 0.100 | 0.0007 | 1.9 |
| Proportional stand movement<br>($R^2 = 0.17$) | Intercept | 1.55167 | 0.348 | <0.0001 | |
|  | area41 | -0.23354 | 0.079 | 0.0030 |  |
| Conifer composition change (%)<br>($R^2 = 0.24$) | Intercept | 0.43100 | 0.074 | <0.0001 | |
|  | Modoc | -0.32900 | 0.107 | 0.0045 |  |
|  | Plumas | -0.13100 | 0.097 | 0.1855 |  |

**Fig 5.**
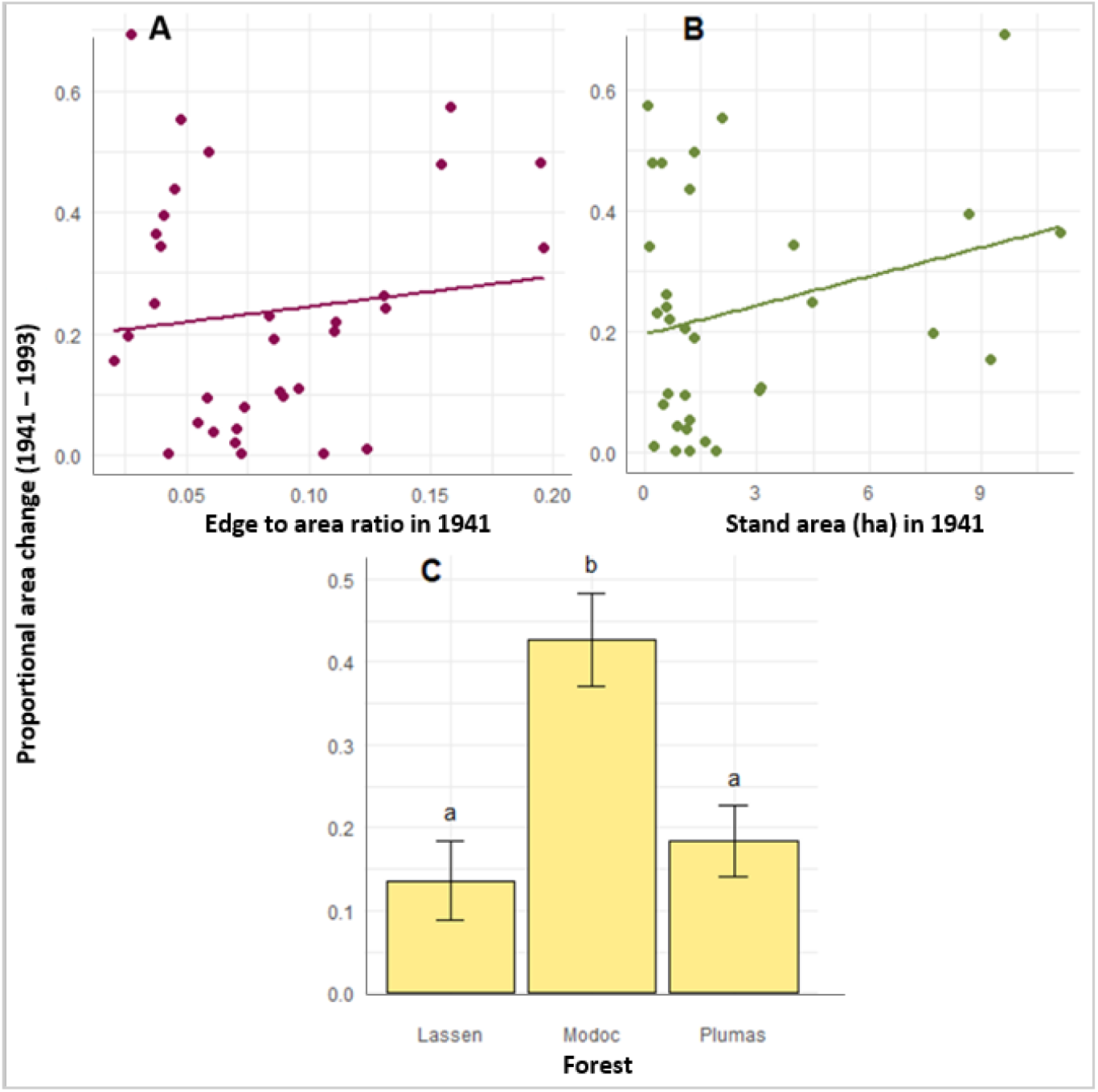
Proportional area change from 1941 to 1993: A) the effect of edge:area ratio in 1941, B) the effect of stand area in 1941 and C) the differences among forests. The plotted dots in A and B are the observed relationship between the two variables, and the lines represent expected area change across range of predictor variable while other variables are held constant at their mean value. The bar graph in C was graphed with pairwise comparison values acquired with emmeans. Error bars represent standard error and letters above each category represent the difference among categories, same letters denoting no significant difference.

While their total area declined in the absence of disturbance, the aspen stands also moved over time and began occupying new areas. During this period of fire exclusion (1941-1993), smaller stands exhibited more movement than larger stands (p = 0.0030) (Fig 6, Table 4).

**Fig 6.**
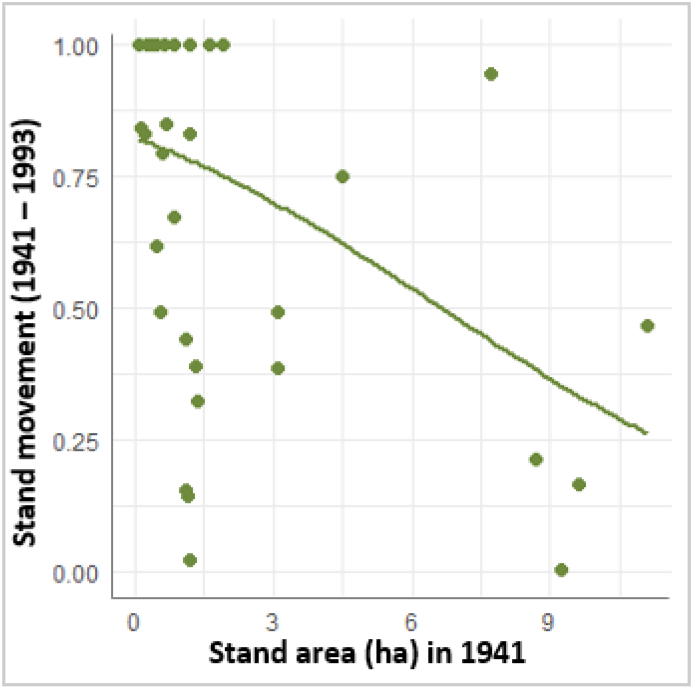
Stand movement from 1941 to 1993 is affected by stand area in 1941. Stand movement on the y-axis represents the portion of a whole stand (1 being the max) that has moved. Plotted dots denote the observed relationship between stand area in 1941 and stand movement between 1941 and 1993. Curve represents expected movement across range of stand areas sampled.

Throughout the period of fire exclusion, conifer composition increased. We did not detect relationships between composition change and any predictor variables except forest. Specifically, conifer encroachment was higher in stands located on the Lassen, followed by the Plumas and lowest on the Modoc (Fig 7, Table 4).

**Fig 7.**
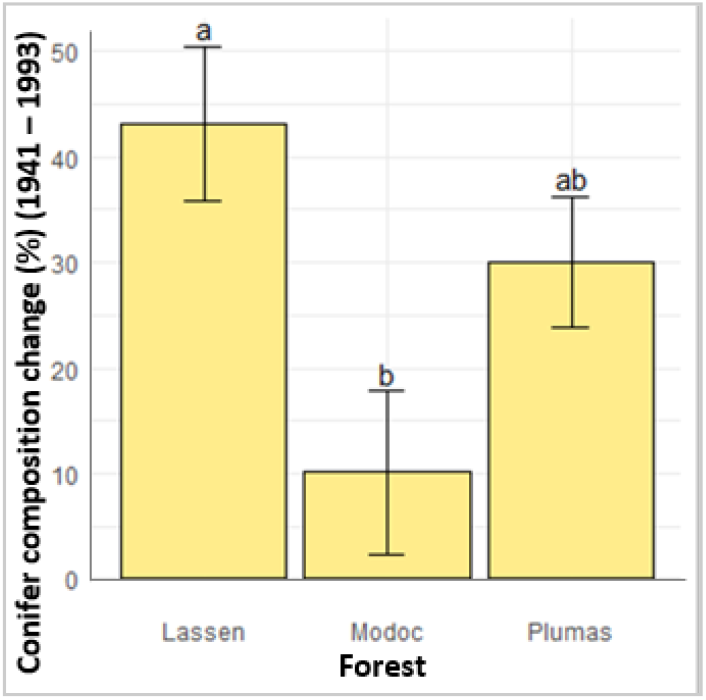
Conifer composition change from 1993 to 1941 differs among forests. Conifer composition on the y-axis shows a change in conifer composition percentage, here all values are positive showing increases among all forests e.g., if a stands conifer composition was 20% in 1941 and 70% in 1993, its’ conifer composition change is 50% (e.g. 70-20 = 50%). Error bars represent standard error of least squares means, and same letter denotes no significant difference among categories.

### Period of Return of Fire (1993 – 2017)

Within the time period where fire returned to the study areas, individual stands that were not exposed to fire continued to decline, while stands that burned expanded in area. Once-burned stands that experienced high severity fire increased in area significantly (p = 0.0045) more than once-burned stands that experience low/moderate severity fire and notably had the greatest increase in area across all fire categories. Area increase fluctuated across edge:area ratio in 1993, but more stands with lower edge:area ratios expanded at greater rates. Greater increases in stand area were also associated with areas that had higher conifer compositions in 1993 (Fig 8, Table 5).

**Table 5.** Models of change over the return-of-fire period (1993-2017) for 33 aspen stands experiencing different fire frequencies and severities: unburned, once-burned experiencing low/moderate severity fire (1burn/LM), once-burned experiencing high severity fire (1burn/H), twice-burned experiencing low/moderate severity fire (2burns/LM) and twice-burned experiencing high severity fire (2burns/H). Variance inflation factor (VIF) quantifies multicollinearity, where a value of 10 or above indicates influential multicollinearity (Kutner et al. 2005).

| Response | Parameter | Coeff | S.E. | Pr(>\|t\|) | VIF |
| --- | --- | --- | --- | --- | --- |
| log(Proportional area change)<br>(Adjusted R<sup>2</sup> = 0.69) | Intercept | -0.876400 | 0.347 | 0.0180 |  |
|  | 1burn/LM | 1.328800 | 0.324 | 0.0003 | 1.6 |
|  | 1burn/H | 2.417500 | 0.441 | <0.0001 | 1.6 |
|  | 2burns/LM | 1.880800 | 0.408 | <0.0001 | 1.6 |
|  | 2burns/H | 1.548100 | 0.420 | 0.0110 | 1.6 |
|  | edge:area93 | -2.212800 | 0.963 | 0.0299 | 1.2 |
|  | %con93 | 1.630800 | 0.507 | 0.0034 | 1.4 |
| Proportional stand movement<br>(R<sup>2</sup> = 0.39) | Intercept | 2.899200 | 0.423 | <0.0001 |  |
|  | area93 | -0.788700 | 0.145 | <0.0001 | 1.4 |
|  | edge:area93 | -15.805300 | 4.601 | 0.0005 | 1.3 |
| Conifer composition change (%)<br>(R<sup>2</sup> = 0.78) | Intercept | 0.005887 | 0.072 | 0.9361 |  |
|  | Modoc | 0.494151 | 0.110 | <0.0001 | 2.5 |
|  | Plumas | 0.074944 | 0.086 | 0.3935 | 2.5 |
|  | FF (1) | -0.426577 | 0.079 | <0.0001 | 1.2 |
|  | FF (2) | -0.503539 | 0.086 | <0.0001 | 1.2 |
|  | riparian | -0.219728 | 0.094 | 0.0284 | 2.6 |
|  | upland | -0.218798 | 0.097 | 0.0332 | 2.6 |

**Fig 8.**
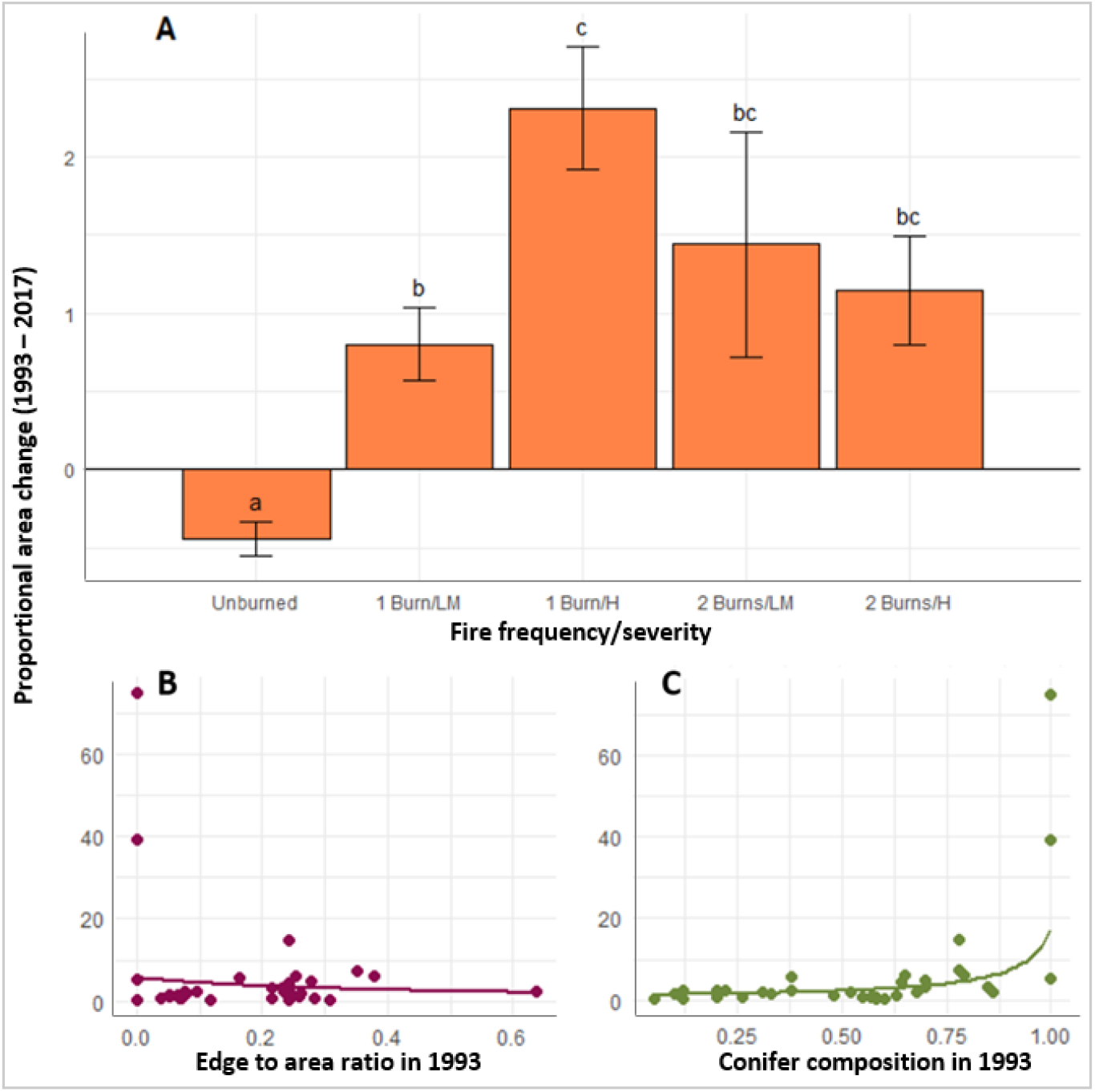
Proportional area change between 1993 and 2017: A) Fire frequency and severity category least squares means and standard errors, B) the effect of edge:area ratio in 1993 and C) the effect of conifer composition in 1993. Error bars represent standard error and letters above each category represent the difference among categories, same letters denoting no significant difference. The plotted dots in B and C are the observed relationships between the two variables, and the curves superimposed over stand data represents expected values across range of predictor variable while other variables are held constant at their mean value.

Aspen stand movement from 1993-2017 followed the same trend as previous years; 379 smaller stands moved significantly (p <0.0001) more than large stands. Aspen stands also moved 380 significantly (p = 0.0005) more when edge:area ratio was lower in 1993 (Fig 9, Table 5).

**Fig 9.**
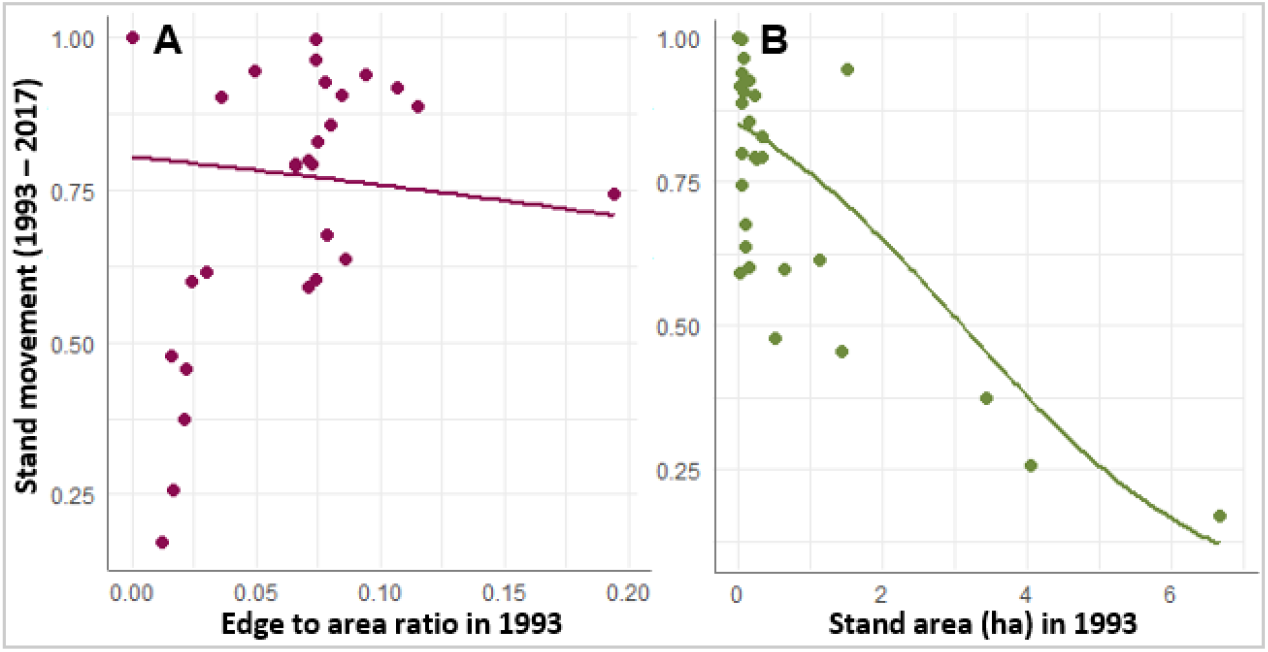
Stand movement from 1993 to 2017: A) the effect of edge to area ratio in 1993 and B) the effect of stand area in 1993. Stand movement on the y-axis represents the proportion of a whole stand area that has moved (1= max.). The curve superimposed over stand data represents expected values across range of predictor variable with the other variable held constant at its mean value.

Conifer composition change was highly influenced by fire; all stands that experienced fire also experienced a significant decrease in conifer composition. Twice-burned stands did not exhibit a statistically detectable difference in decline of conifer composition than once-burned stands. After accounting for the influences of disturbance type and site type in the regression analysis, stands on the Modoc experienced a small average increase in conifer composition while the Lassen and Plumas stands experienced large decreases in conifer composition. Pure riparian and upland stands both showed significantly (p = 0.0284, 0.0332) more conifer loss than mixed type stands but did not differ significantly (p = 1.0) from each other (Fig 10, Table 5).

**Fig 10.**
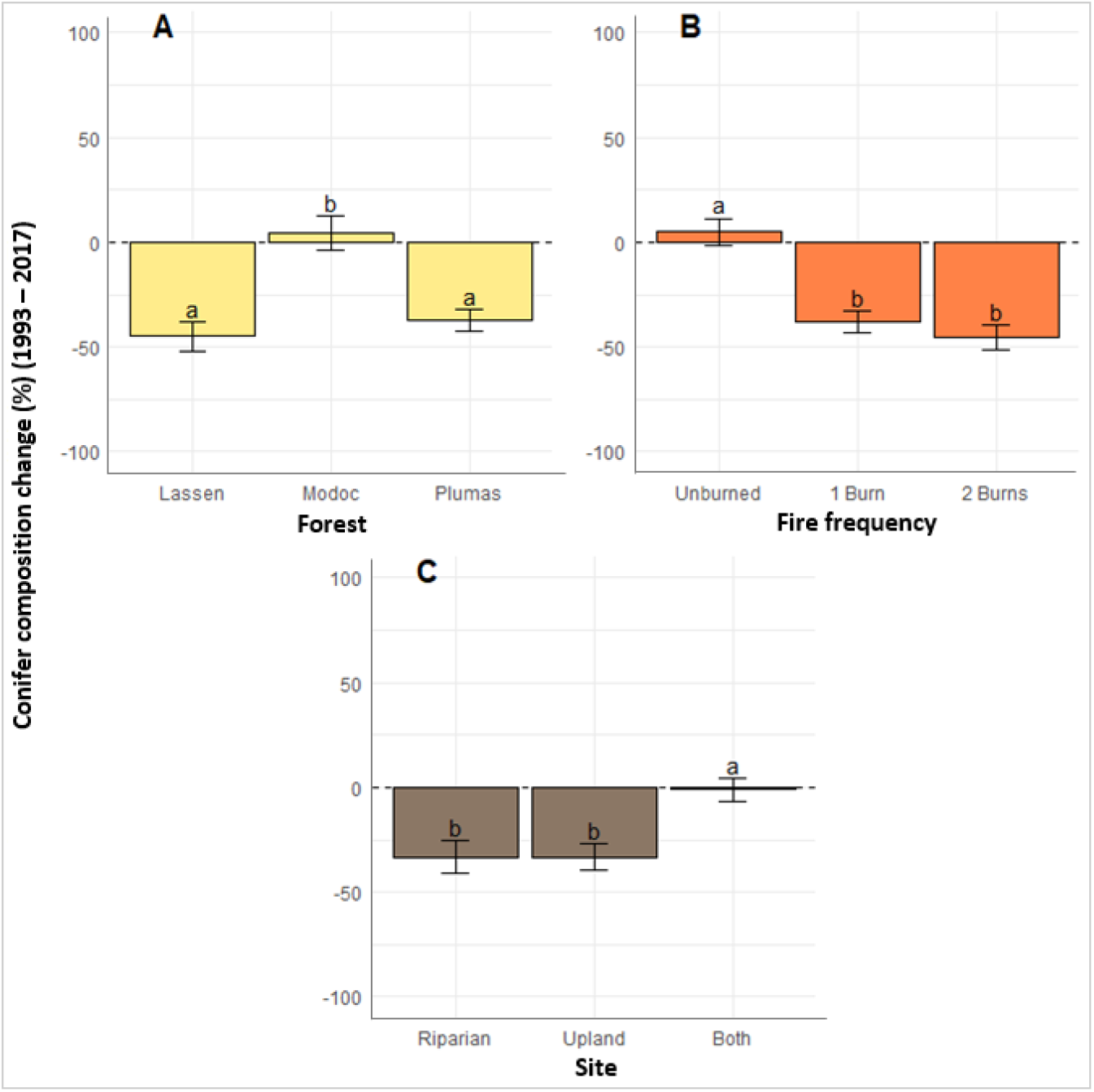
Conifer composition change between 1993 and 2017: A) differences among forest, B) fire frequencies and C) site types, e.g. if a stand’s conifer composition was 50% in 1993 and 10% in 2017, it’s conifer composition change was −40%. Graphed values are from pairwise comparisons acquired with emmeans. Error bars represent standard error. Categories with same letter were not significantly different.

## Discussion

### The Role of Wildfires in Declining and Expanding Aspen Stands

We observed spatial changes in aspen stands in the absence and presence of wildfire. Without fire on the landscape, aspen stand area declined across all stands and on all forests studied. During our study period of 52 years without wildfire, aspen stand area declined by 76% on average. Four of our 32 studied stands were completely undetectable in 1993 aerial photography. Di Orio et al. [34] observed aspen decline on the Modoc National Forest, also within a similar time range (1946-1994), finding a 24% decline in total area studied. This was lower than our observations of decline on the Modoc which had the least decline of our three study areas at about 54% (1941-1993). Aspen on the Lassen and the Plumas National Forests had higher rates of decline (90% and 73%, respectively) over the same 52-year period. In a study of aspen in the western U.S., Bartos [12] reported on a century of decline which ranged from 49% to 95%.

The decline of aspen appeared to be linked to the process of succession to conifers within these aspen-conifer stands. In the absence of fire, conifer composition increased by 29% on average over 52 years. This is consistent with 50-year data for Lassen Volcanic National Park in northeastern California, where aspen stands deemed to be undergoing rapid succession to conifer had 29% less aspen cover and 46% greater conifer cover at the end of their study period [51].

The steady establishment of conifers within the aspen stand footprints is assumed to be correlated to lack of disturbance. Our study observed a significant increase in conifer composition during the period when fire was absent, and a significant decrease in conifer composition when fire was present. This suggests that the lack of fire or other disturbance, which remove conifers and release aspen from competition is a driving cause for aspen decline in these areas. Restoration treatments found to promote aspen regeneration and growth include conifer removal alone [52, 37] or combined with prescribed fire [23]. Expansion between 1993 and 2017 was also affected by the percentage of conifers present in 1993 prior to burning. During the return-of-fire period, burned stands experienced greater expansion where conifer composition was higher at the beginning of the period. These stands may have exhibited a greater response from the removal of conifer competition as well as stimulation of regeneration by fire [53], in turn resulting in a greater rate of area expansion over our study period.

In the presence of fire, aspen stands expanded and reoccupied areas of former stand footprints and new areas. Total stand area that experienced a single fire event within the 24-year period of return of fire increased in stand area by 60% from total area in 1993. Total stand area that burned twice between 1993 and 2017 increased in stand area by 75%. Even after expansion, these new areas were still only a fraction of observed stand sizes in 1941. It is uncertain whether expansion will continue in the future, and if compound disturbances are more beneficial than infrequent disturbances. We recommend studying a larger sample size of aspen stands in light of the variability exhibited by our data for stand area expansion, stand movement, and percent conifer composition.

Expansion where fire has occurred may also be a consequence of the removal of conifers due to high severity fire. Stands that burned at low or moderate severity increased in stand area by 52% from 1993 to 2017. Stands that burned at high severity increased in area by 82%. A major consequence of fire severity is conifer removal, as high severity stands were classified as having 100% conifer mortality within the period of return of fire. Fire severity also correlates with enhanced re-sprouting of other vegetation such as black oak (*Quercus kelloggii*) in this forest type [54], especially after reburn [55, 56, 57]. In the absence of high-severity fire, heavy or frequent conifer removal will be needed to restore aspen dominance in stands undergoing succession from aspen to conifer [58].

We also observed greater area expansion and movement in aspen stands that had lower edge:area ratios. Where large populations of ungulate grazers are present, aspen stands with greater edge are less likely to successfully regenerate [59]. Greater edge was more commonly observed in aspen stands interspersed with conifers. Aspen regeneration and recruitment decreases when more conifers are present in a stand [60, 29]. The greater movement of small aspen stands with lower edge:area ratio compared to larger stands may indicate that smaller stands are more susceptible to replacement by conifers if migration into an adjacent area is impeded by dense conifer forest or other obstacles.

When using aerial photo sets from different years, with different quality and resolution, we encountered a problem with a confounding of time and technology. With the improvement of technology throughout time, photo quality increased progressively. Because of this, decisions on whether or not to delineate areas of aspen had to be consistent with what was visible within the oldest photos (1941), even though much younger trees were observable in the most recent imagery. This resulted in the exclusion of small, young individuals in all years to avoid bias towards including more aspen area in later years that may only be detectable due to better photo quality. We were also limited by choosing stands that were present at the time of recent field delineation, as we used these recent GPS polygons to locate areas of aspen in the historical imagery. This may have prevented us from observing stands that were present in previous years and had since been lost from our study areas within our study period.

Another limitation appeared when trying to isolate the effect of time since fire had burned a stand in our models for the period of return of fire. Including the number of years since fire as a predictor variable was not possible while trying to compare to the unburned stands where the last year they had burned was unknown. Because of this, we expect that there may be an unexplained effect of date-of-fire from the forest where stands were located. We also may have underestimated responses in some twice-burned stands because each forest had different years of fire within our study period. The longest time between the latest fire and the aerial imagery date was on the Modoc (16 years), and the shortest time since fire was on the Lassen (5 years). Therefore, twice-burned aspen regenerating after fire on the Lassen had fewer growing seasons to respond to the effect of fire and would not be as large or visible after the most recent fire. Field assessment of the same aspen stands on the Lassen revealed many areas with small new root suckers that were not included in aspen stands delineated by remote sensing. If the Lassen stand areas had included these suckers, their post-fire area expansion may have been even greater.

### Implications for Management

All aspen stands that burned appear to be expanding in area post-fire. However, the sustainability of this upward trend is unknown. Monitoring these stands into the future would be required to fully understand how often these stands need to burn to maintain or expand their area into the future. The continued decline of stands that did not burn during the entire study period (1941-2017) suggests that these stands may be lost without active restoration. Although our study has shown how wildfire disturbances favor aspen over conifer, fire may not be the most feasible restoration tool to use. For aspen stands that are nearing complete disappearance, prescribed fire or fire surrogates such as mechanical removal of conifers, may provide an immediate benefit needed to promote aspen regeneration and growth by reallocating growing space to aspen. Regeneration from suckering may be promoted from options other than fire as well, including manual breakage of the roots [10]. In areas where aspen is present and wildfires are expected in the future, land managers should have wildland fire use plans in place in preparation for the opportunity to introduce natural fire into aspen stands. If declining stands are not exposed to fire or fire surrogates, the effects of succession to conifers may lead to complete overtopping of aspen and loss of entire aspen stands. Conversely, the clear benefits of wildfire to aspen stands in our study highlights the potential for fire to reverse aspen decline in this region, even after a long period of fire exclusion.

